# Corpusome, a cross-body-site human microbiome corpus for representation learning

**DOI:** 10.64898/2026.08.28.747922

**Authors:** Hao Xuan, Yu Huang, Jiang Bian

## Abstract

Machine-learning models of the human microbiome are trained mostly on stool samples from single cohorts, limiting cross-body-site representation and cross-study generalization. Progress is constrained less by algorithms than by the absence of a harmonized multi-body-site corpus carrying the technical metadata needed to model, rather than ignore, batch structure. Here we release Corpusome, a harmonized two-tier cross-body-site human microbiome corpus for representation learning: a harmonized corpus of 187,546 human microbiome samples integrating standardized profiles from curatedMetagenomicData, the American Gut Project, and the EBI MGnify platform. Corpusome follows a two-tier design preserving both functional depth and cross-body-site breadth: a shotgun tier (22,588 samples, 93 studies) with species- and pathway-level profiles, and a 16S tier (164,958 samples, from a full pull of 708 MGnify studies) with genus-level profiles extending coverage to oral, skin, respiratory, and urogenital sites. It spans six body sites and two modalities, with harmonized metadata for batch-aware modelling. Body-site signal exceeds technical/source variance in the 16S tier by approximately 2.4-fold. Corpusome is freely available at https://xuan13hao.github.io/Corpusome/.

## Background & Summary

The human microbiome varies profoundly across body sites: the communities of the gut, mouth, skin, airways, and urogenital tract differ in composition, diversity, and function. Yet the datasets used to train computational models of the microbiome are dominated by a single niche (stool) drawn from individual studies and are typically confined to one sequencing modality. Models trained this way learn features that are entangled with cohort-specific technical artefacts and that do not transfer to other body sites or studies. Self-supervised “foundation model” approaches to microbial data have made this limitation explicit, identifying multi-body-site integration and cross-study generalization as the principal open problems.

Addressing those problems requires training data with three properties that no single public resource provides at once: (1) breadth across body sites, (2) depth of functional characterization, and (3) rich, harmonized technical metadata that lets biological and batch effects be separated during modelling. curatedMetagenomicData (cMD) [1] offers depth, uniformly reprocessed shotgun taxonomic and functional profiles for tens of thousands of samples, but is dominated by stool. The American Gut Project (AGP) [2] contributes a large, fully public citizen-science cohort with genuine within-subject multi-site sampling, but only as 16S amplicon data. The EBI MGnify platform [3] hosts standardized profiles for hundreds of thousands of samples across every human body site, in both modalities, but serves them per-sample through a rate-limited API rather than as ready analysis matrices.

Here we integrate these three resources into Corpusome, a single harmonized corpus. Rather than force incompatible data types into one matrix, which would confound biological signal with primer-region and profiler differences, we adopt a two-tier design that keeps shotgun and 16S profiles as linked-but-separate feature spaces, with modality recorded as an explicit metadata field. The result is Corpusome: 187,546 samples spanning six body sites and two modalities, with per-sample body site, study/source, sequencing platform, and host covariates retained for batch-aware modelling. We provide feature tables (species, pathways, genera), a unified sample metadata catalogue, and a machine-readable data-records inventory. Corpusome is designed as a pretraining and benchmarking resource; disease-labelled cohorts for downstream fine-tuning are described as external add-ons and are not part of this release.

## Methods

Corpusome is built from three public resources through a sequence of acquisition, harmonization, and validation steps, described below (Figure 1).

**Figure 1.**
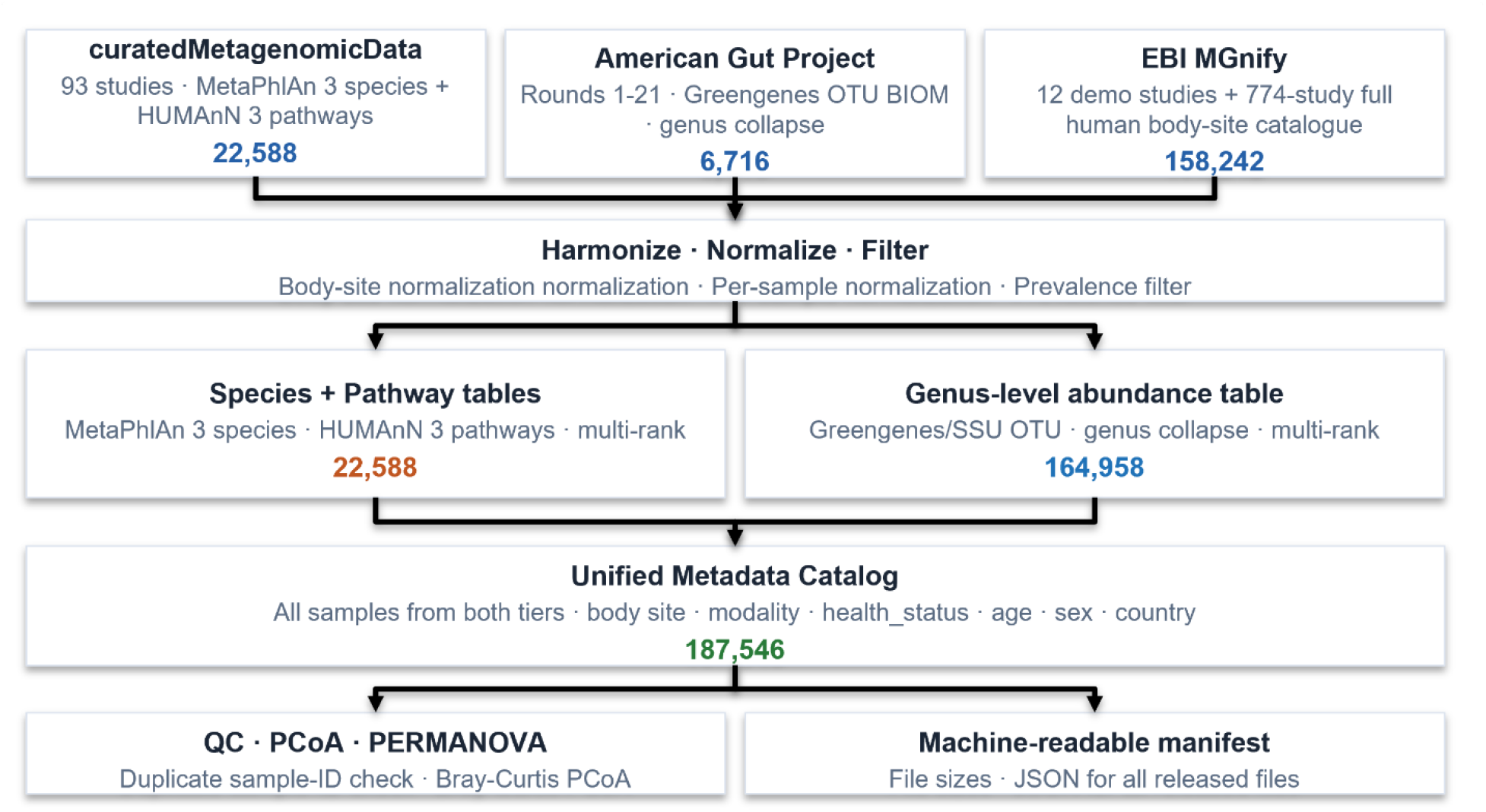
Construction pipeline for Corpusome. Standardized profiles from three public resources, curatedMetagenomicData, the American Gut Project, and EBI MGnify, are harmonized, normalized, and prevalence-filtered into two tier-specific feature tables: a shotgun species/pathway table and a 16S genus table. Sample metadata from all sources are merged into a unified catalogue of 187,546 samples, followed by technical validation (PCoA/PERMANOVA) and a machine-readable file manifest.

### Shotgun tier: curatedMetagenomicData

Standardized shotgun profiles were obtained from the curatedMetagenomicData resource, accessed through its current Bioconductor release. Taxonomic profiles are MetaPhlAn 3.0 [4, 5] clade relative abundances; functional profiles are HUMAnN 3.0 pathway abundances[5, 6]. Because the underlying data releases accumulate by study-addition date rather than as a single cumulative snapshot, complete coverage of all 93 studies required combining multiple releases, using the most comprehensive release as the primary source and earlier releases to fill in individual studies it did not yet cover.

Species-level taxonomic assignments were retained (excluding strain-level resolution) and assembled into a samples-by-species abundance matrix. Functional profiles were retained at the level of community-wide metabolic pathways, excluding species-specific breakdowns, while still capturing the fraction of reads that could not be mapped to a known gene or integrated into a named pathway. Because raw sample identifiers are not unique across studies, each sample was assigned a composite identifier combining its study and original sample label. All 93 studies were retrieved successfully, with no download failures.

### 16S tier: American Gut Project

The AGP precomputed release was obtained from the project’s public data mirror (open access; ENA study PRJEB11419 /ERP012803) [2]. We used the standard closed-reference OTU tables provided with the release, generated against the Greengenes 13_8 reference database [7] from 16S rRNA V4-region amplicons (515F/806R primers). OTU taxonomy was collapsed to the genus level by summing counts within each full lineage; genera unresolved at that rank were labelled by their deepest assigned rank. AGP provides genuine within-subject multi-site coverage: 363 participants were sampled at more than one body site.

### 16S tier: MGnify full pull

To extend 16S coverage across all human body sites at scale, we performed a full pull of the MGnify human body-site catalogue3 (774 catalogued studies, of which 770 were included in the pull). Rather than querying MGnify’s rate-limited per-sample interface, we retrieved each study’s combined SSU rRNA taxonomy table, a single summary covering all of that study’s samples with the full taxonomic lineage, which reduced acquisition time roughly twelve-fold and made a whole-catalogue pull feasible. Of the 770 studies, 708 provided a usable combined table, yielding 157,915 samples; the remaining 62 had no such table (being fungal-only or shotgun-only studies) and are logged for provenance. Lineage information was aggregated to each taxonomic rank from phylum through genus.

### Harmonization

Body-site labels from all sources were mapped onto a controlled vocabulary spanning stool, oral, skin, respiratory, urogenital, and milk. Because the MGnify full pull uses a different reference taxonomy (SILVA) [8] from the American Gut Project (Greengenes), the 16S genus-level tables were harmonized by matching taxa on their genus-level name; of the 683 genera present in the earlier 16S tier, 514 are shared with the MGnify vocabulary. Each feature table was converted to per-sample relative abundance and filtered to remove rare, uninformative taxa or pathways, retaining only those present in a meaningful minimum share of samples. The expanded 16S genus table spans 164,958 samples across 977 genera; the shotgun tier retains 713 species and 510 pathways. Coarser taxonomic ranks, from phylum through family, are released alongside the finest resolution for both tiers. Sample-level metadata from all sources were merged into a single catalogue capturing, for every sample, its body site, sequencing modality, study or data source, geographic region, processing pipeline, host age and sex, subject identifier, country, sequencing platform, and any associated disease or health status.

### Data Records

Corpusome is deposited as a single versioned archive (see Data Availability). Table 1 lists the released records; the complete schema, per-file checksums, and provenance are documented in an accompanying machine-readable inventory. Figure 2 summarizes total sample counts by body site, combining both tiers (see also Table 2). Figure 3A breaks this down by sequencing modality (shotgun vs 16S).

**Table 1:**
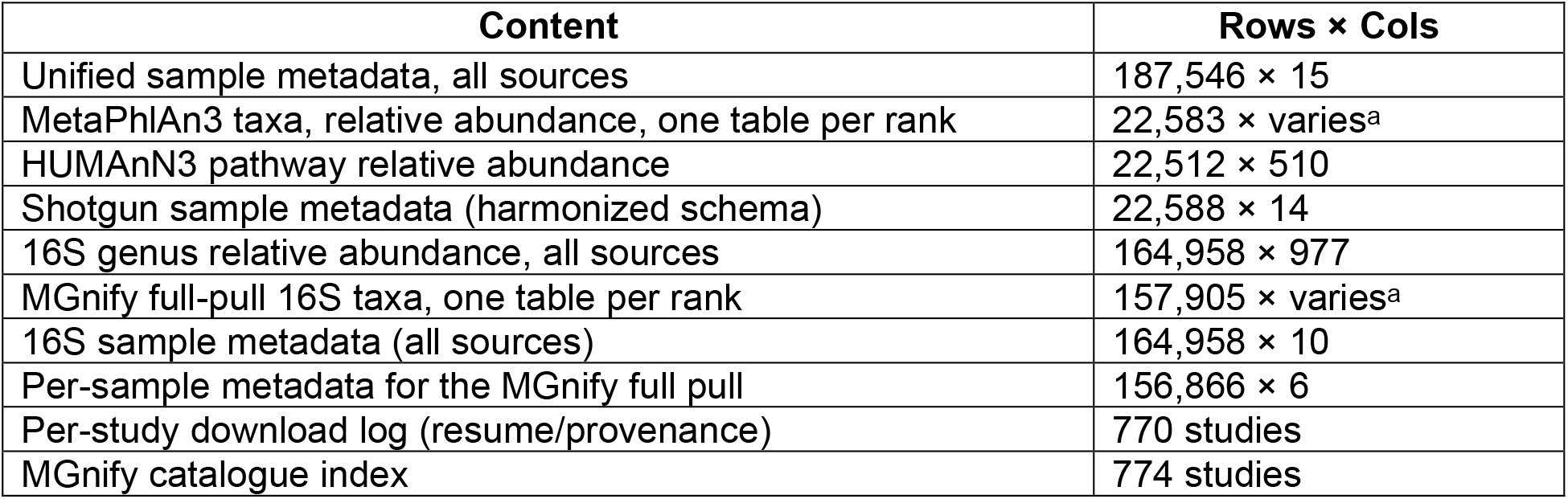
Released data records. ^a^ Column count varies by taxonomic rank (phylum through species/genus); exact per-rank dimensions are given in the accompanying data inventory.

**Table 2:**
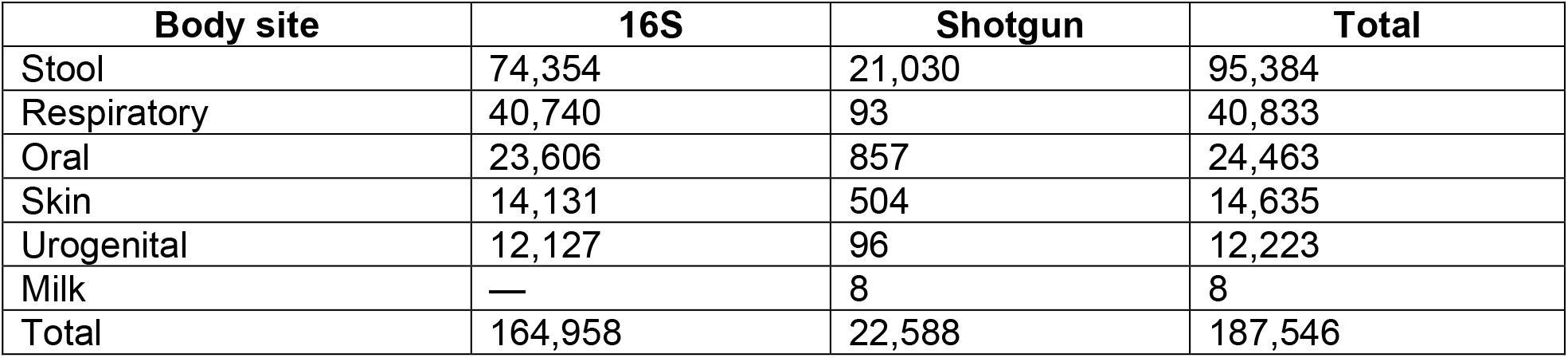
Corpus composition by body site and modality (sample counts).

| **Body site** | **16S** | **Shotgun** | **Total** |
| --- | --- | --- | --- |
| Stool | 74,354 | 21,030 | 95,384 |
| Respiratory | 40,740 | 93 | 40,833 |
| Oral | 23,606 | 857 | 24,463 |
| Skin | 14,131 | 504 | 14,635 |
| Urogenital | 12,127 | 96 | 12,223 |
| Milk | — | 8 | 8 |
| Total | 164,958 | 22,588 | 187,546 |

**Figure 2.**
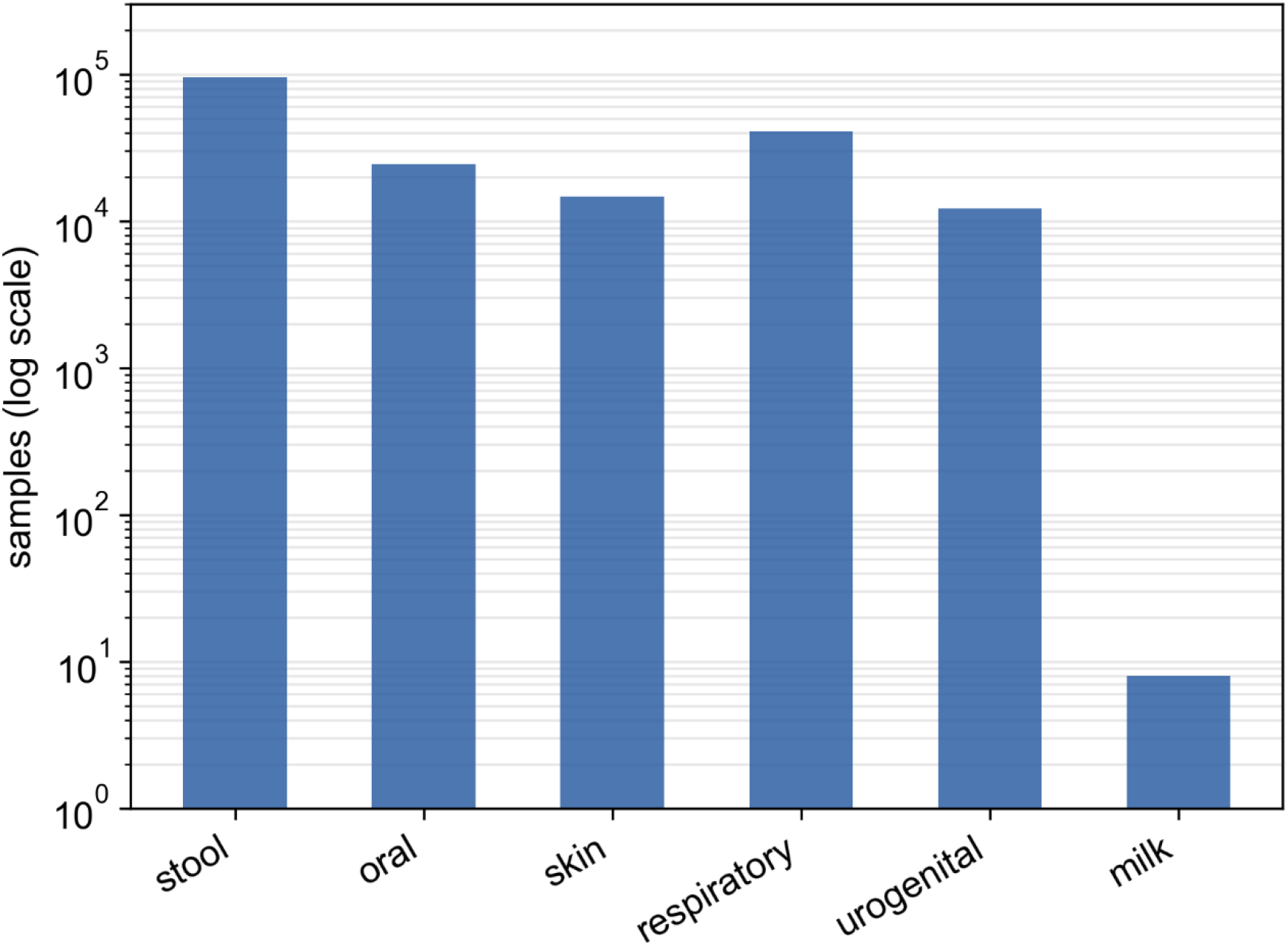
Total sample counts by body site (16S and shotgun tiers combined), log scale. Stool is the largest category (95,384 samples), followed by respiratory (40,833), oral (24,463), skin (14,635), and urogenital (12,223); milk is a minor category (8 samples).

**Figure 3.**
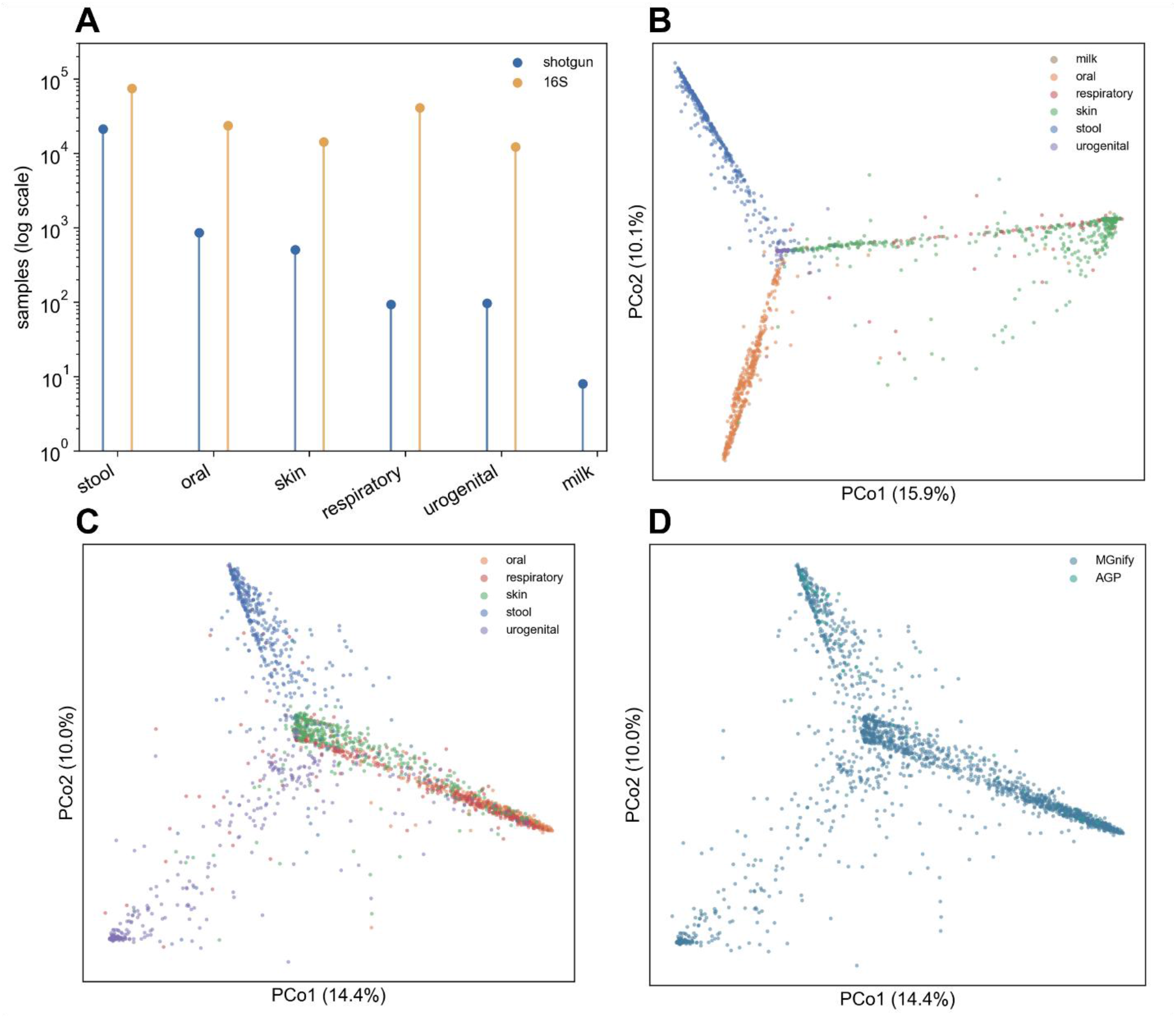
Technical validation of the harmonized corpus. (**A**) Sample counts by body site and sequencing modality, log scale (shotgun, blue; 16S, orange). (**B**) PCoA of Bray–Curtis dissimilarities among shotgun-tier species profiles, colored by body site. (**C**) PCoA of 16S-tier genus profiles, colored by body site. (**D**) The same 16S-tier ordination as in (**C**), colored by data source (AGP vs MGnify).

#### Taxonomic resolution

To support modelling at multiple taxonomic scales, each tier is released at every rank from phylum to its finest available resolution — species for the shotgun tier (MetaPhlAn3), genus for the 16S tier (Greengenes lineages), plus phylum/class/order/family for both. Each rank is a separate per-sample relative-abundance table, prevalence-filtered under the same rule, and derived by summing the finest-rank profiles within each parent lineage (rank tables therefore nest consistently: a phylum abundance equals the sum of its constituent genera/species). Users select the resolution appropriate to their model; coarser ranks are denser and less study-specific, finer ranks carry more biological detail.

#### Health status

Each sample is annotated in the metadata catalogue as healthy, diseased, or unlabeled. Corpusome is predominantly but not exclusively healthy: within the shotgun tier, 14,566 samples (64.5%) are from healthy/control participants and 8,022 (35.5%) carry a disease label spanning 141 conditions (most frequently inflammatory bowel disease, type 2 diabetes, and colorectal cancer). The 16S tier (American Gut Project, MGnify) is not disease-annotated and is marked unlabeled (164,958 samples). This healthy-plus-perturbed mixture is deliberate for a pretraining corpus: it exposes a model to both baseline and altered community states. It is distinct from the dedicated disease cohorts used for downstream fine-tuning, which are out of scope here (see Usage Notes). Every feature table is linked to the metadata catalogue through a shared sample identifier, and all feature-table values are expressed as per-sample relative abundances that sum to one.

### Technical Validation

#### Body site is the dominant axis of variation

Principal-coordinates analysis on Bray–Curtis dissimilarities separates samples by body site in both tiers (Figure 3B, C). We quantified this with a PERMANOVA-style variance partition (the ratio of between-group to total sum of squares) on body-site-stratified subsamples. In the 16S tier, body site explains *R*^2^ = 0.32 of community variation versus *R*^*2*^ = 0.13 for data source, biological signal exceeds the technical/source effect by 2.4× (Figure 3D). In the shotgun tier, body site (*R*^2^ = 0.28) and study (*R*^2^ = 0.31) explain comparable variance; because cMD studies differ systematically by body site, these two effects are partially confounded and should be modelled jointly rather than interpreted as independent.

#### Integrity and completeness

No duplicate sample identifiers occur across the 187,546 samples. Body site is labelled for 100% of samples; host age, sex, and subject identifiers are present for 81.6%, 88.6%, and 100% of samples, respectively; the 16S tier is dominated by the MGnify full pull (156,866 of 164,958 16S samples), which was deposited without matched host age, sex, or subject-identifier fields, so these covariates should be read at the tier level rather than pooled across the full corpus. The American Gut Project’s within-subject design is preserved: 363 participants are sampled at more than one body site, supporting paired cross-site analyses.

#### Reproducibility

The released pipeline reconstructs every Corpusome table from primary sources; running the harmonization and validation stages on the acquired inputs reproduces the reported dimensions and validation statistics exactly. Per-file SHA-256 checksums are provided.

### Usage Notes

The shotgun (species/pathway) and 16S (genus) tiers use different profilers and reference taxonomies and are provided as linked-but-separate feature spaces. Modality is an explicit metadata column and should be used as a conditioning or batch variable, not marginalized away by naïvely concatenating the tiers. Study (shotgun) and source (16S) are recorded per sample. Given the confounding between study and body site in the shotgun tier, we recommend conditioning on study identity (or applying reference-based normalization) and reporting results with and without batch correction.

Stool dominates both tiers; non-stool sites are comparatively better represented in the 16S tier. For balanced pretraining, consider body-site-stratified sampling, as used for the validation ordinations. The 16S tier already incorporates the full MGnify human body-site pull (708 studies). The acquisition step is resumable and tracks per-study status, so re-running it picks up any studies that failed or were newly added to MGnify; the shotgun tier can likewise be grown from newer curatedMetagenomicData releases.

## Data Availability

Derived feature tables, the metadata catalogue, the machine-readable inventory, and validation outputs are deposited under a Zenodo (https://doi.org/10.5281/zenodo.22150633). Raw sequence data remain in their original public archives and are cited rather than re-deposited: per-study ENA/SRA accessions for the shotgun tier (recorded in the metadata), ENA PRJEB11419/Qiita study 10317 for the American Gut Project, and the MGnify study accessions listed in the accompanying download manifest.

## Code Availability

All code, reproducible pipeline scripts, containerized software environment specifications, and extended documentation are permanently archived in Zenodo (https://doi.org/10.5281/zenodo.22150633).

## Acknowledgements

We are grateful to the curatedMetagenomicData, American Gut Project, and MGnify teams for generating and publicly releasing the data underlying this resource.

## Author Contributions

HX performed data acquisition, harmonization, and technical validation. HY and JB supervised the study. The original draft was written by HX, and the final manuscript was reviewed and edited by HX, YH, and JB.

## References

1. Pasolli, E., et al., Accessible, curated metagenomic data through ExperimentHub. Nat Methods, 2017. 14(11): p. 1023–1024.

2. McDonald, D., et al., American Gut: an Open Platform for Citizen Science Microbiome Research. mSystems, 2018. 3(3).

3. Richardson, L., et al., MGnify: the microbiome sequence data analysis resource in 2023. Nucleic Acids Res, 2023. 51(D1): p. D753–D759.

4. Blanco-Miguez, A., et al., Extending and improving metagenomic taxonomic profiling with uncharacterized species using MetaPhlAn 4. Nat Biotechnol, 2023. 41(11): p. 1633–1644.

5. Beghini, F., et al., Integrating taxonomic, functional, and strain-level profiling of diverse microbial communities with bioBakery 3. Elife, 2021. 10.

6. Jung, S., et al., Building resource-efficient community databases using open-source software. Database (Oxford), 2025. 2025.

7. McDonald, D., et al., Greengenes2 unifies microbial data in a single reference tree. Nat Biotechnol, 2024. 42(5): p. 715–718.

8. Chuvochina, M., et al., SILVA in 2026: a global core biodata resource for rRNA within the DSMZ digital diversity. Nucleic Acids Res, 2026. 54(D1):p. D334–D341.

